# Action Potentials and Na^+^ voltage-gated ion channels in Placozoa

**DOI:** 10.1101/2020.08.09.243113

**Authors:** Daria Y. Romanova, Ivan V. Smirnov, Mikhail A. Nikitin, Andrea B. Kohn, Alisa I. Borman, Alexey Y. Malyshev, Pavel M. Balaban, Leonid L. Moroz

## Abstract

Placozoa are small disc-shaped animals, representing the simplest known, possibly ancestral, organization of free-living animals. With only six morphological distinct cell types, without any recognized neurons or muscle, placozoans exhibit fast effector reactions and complex behaviors. However, little is known about electrogenic mechanisms in these animals. Here, we showed the presence of rapid action potentials in four species of placozoans (*Trichoplax adhaerens* [H1 haplotype], *Trichoplax sp*.[H2], *Hoilungia hongkongensis* [H13], and *Hoilungia sp*. [H4]). These action potentials are sodium-dependent and can be inducible. The molecular analysis suggests the presence of 5-7 different types of voltage-gated sodium channels, which showed substantial evolutionary radiation compared to many other metazoans. Such unexpected diversity of sodium channels in early-branched animal lineages reflect both duplication events and parallel evolution of unique behavioral integration in these nerveless animals.

**Highlights:**

- Placozoans are the simplest known animals without recognized neurons and muscles
- With only six morphological cell types, placozoans showed complex & rapid behaviors
- Sodium-dependent action potentials have been discovered in intact animals
- Voltage-gated sodium channels (Na_v_) in Placozoa support a rapid behavioral integration
- Placozoans have more Na_v_ channels that any studied invertebrate animal so far
- Diversification of Na_v_-channels highlight the unique evolution of these nerveless animals

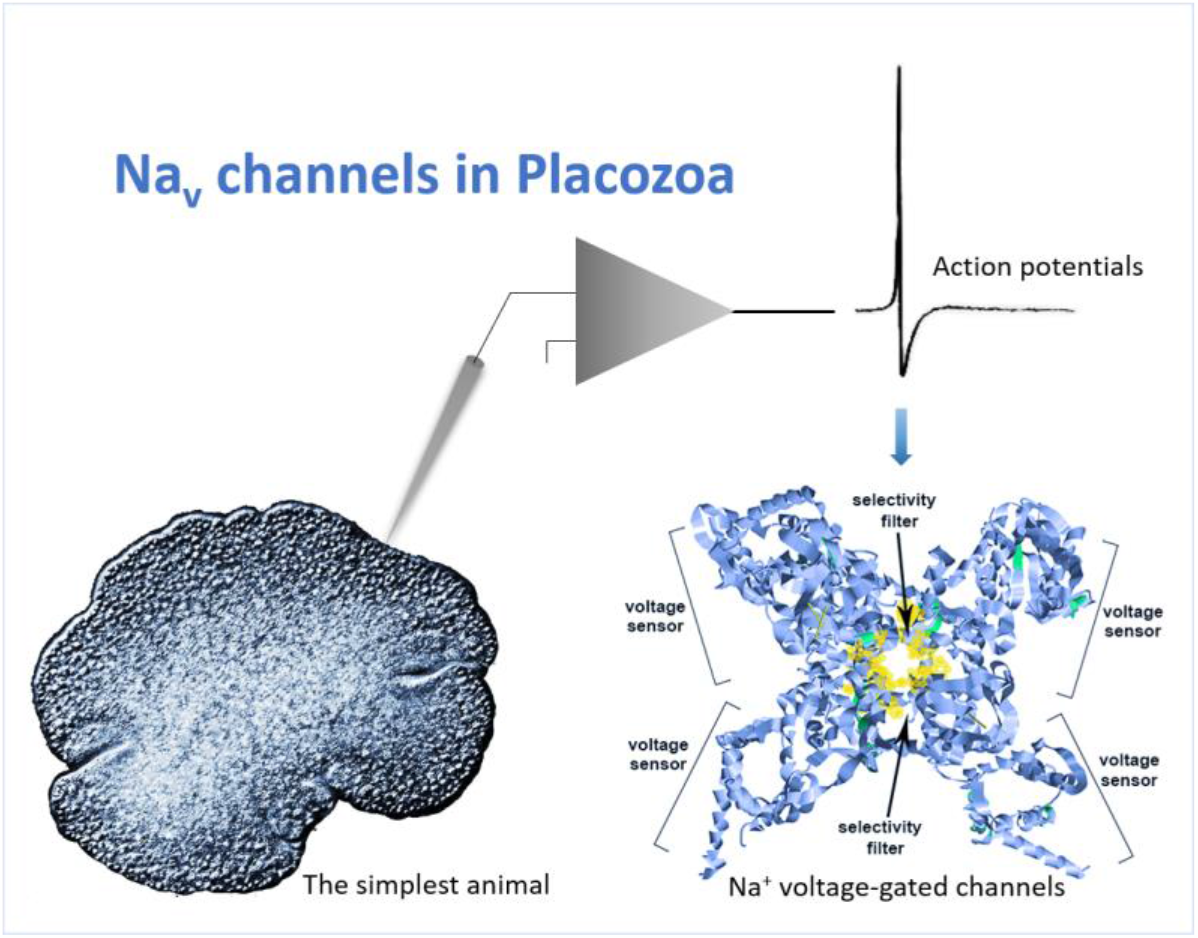

## 1. Introduction

The origin and early evolution of electrogenic mechanisms for behavioral integration in animals are elusive[1–3]. A diverse set of voltage-gated ion channels and all-or-none action potentials have been described in ctenophores - descendants of the earliest-branching metazoans, which independently evolved neuro-muscular systems[4,5]. On the other hand, Porifera might exemplify secondary simplification of features, and many sponges lost some ancestral traits such as sodium voltage-gated channels[6,7]. However, slow (likely calcium) action potentials can be recorded from glass sponges[8,9].

Thus, representatives of Placozoa (the third branch on the animal tree of life) can be critical reference models to decipher the development of integrative mechanisms in Metazoa[10–14]. Although placozoans might contain more than one hundred marine species[15], most information about the group is based on the study of *Trichoplax adhaerens -* the founding member of the phylum. These 0.5-2 mm disc-shaped benthic grazers are the simplest free-living animals[16], with only six morphologically recognized cell types[17].

Despite stunning morphological simplicity, placozoans have a complex behavioral repertoire[18–23], including social behaviors[24] and ultrafast contractions[25] as well as a complex system of intercellular signaling[17,18,22,26–30]. But the cellular bases of behavior in *T.adhaerens* and related species are unknown. We hypothesize that placozoans have developed all-or-none action potentials to support the rapid propagation and integration of electrical and chemical signals across cell layers. A specialized meshwork of fiber cells, located in the middle layer of cells, was considered to be an analog of the neural and muscular systems[16], but no synapses and no gap junctions have been described morphologically[16,17]. The innexins and connexins – the canonical gap junction proteins are not encoded in the sequenced genome of *T.adhaerens* and its kin[14,31,32]. Adherent junctions do facilitate diffusion of potential nutrients into the animals[33], but it is unknown if they participate in the propagation of any electrical signals.

Ultrasmall sizes of most of the placozoan cells (3-5μm), their extremely fragile nature, significantly limit the application of patch-clamp protocols for direct electrophysiological studies. Furthermore, a very high level of autofluorescence also restricts the usage of voltage-sensitive dyes for these animals. Both voltage-gated T-type calcium[34] and leak sodium[35] ion channels have been recently cloned and expressed in heterologous systems. These studies provided the first insights into the placozoan electrophysiology. Also, two types of sodium voltage-gated ion channels have been indentified in *Trichoplax*[6,36,37], and they might belong to the Na_v_2-like family of the channels with possible Ca^2+^- selectivity[37–40]. However, no successful recordings were performed from intact animals, and it is unclear whether regenerative and/or Na^+^-dependent action potentials exist in placozoans.

Here, we performed electrophysiological tests on representatives of two placozoa genera (*Trichoplax* and *Hoilungia*) and provided the evidence for the presence of rapid Na^+^-dependent action potentials. We also revealed a surprising diversity of sodium voltage-gated channels in all four investigated species as well as candidates for other diverse families of cationic channels. Our data suggest parallel evolution and prominent diversification of mechanisms controlling excitability in these nerveless animals.

## 2. Materials and methods

### 2.1. Animals

Three different species, *Trichoplax adhaerens* (haplotype H1), *Trichoplax* sp.(H2), and *Hoilungia hongkongensis*(H13)[31], were maintained in the laboratory culture; animals were fed on rice grains and algae as described elsewhere[23]. *Hoilungia* sp. (H4) were maintained in 10-40 L marine aquaria and fed on three species of algae (*Tetraselmis marina*, *Spirulina versicolor*, *Leptolyngbya ectocarpi*)[28,41].

### 2.2. Electrophysiological experiments

In all tests, animals were placed in 3mm Petry dishes, and experiments were conducted under differential contrast microscopy (DIC, Olympus BX51WI microscope, n=49 specimens). Extracellular recordings were performed using glass microelectrodes. The microelectrodes were made from borosilicate glass capillary (BF150-86-10) on a Sutter p-1000 puller (Sutter,USA) and filled with artificial seawater or equimolar sodium-free solution. The extracellular signals were filtered from 300 to 10000 Hz, amplified (x50) with MultiClamp 700B amplifier (Molecular Devices,USA), digitized at 20 kHz using Digidata 1500 and recorded using PCLAMP software (Molecular Devices,USA).

To reduce movements, placozoans were immobilized in 1% agarose made either artificial seawater (ASW: NaCl- 450mM, KCl-13.4mM, MgCl_2_-24mM, CaCl_2_-9.5mM, MgSO_4_–5mM; pH=8.0) or sodium-free seawater, where NaCl was replaced by *N*-methyl-d-glucamine (NMDG, 390mM, pH=8.0)[42]. Initially, 1% agarose and seawater were heated to 40oC, and then we placed a small (<0.1-0.2 mL) drop of the agarose solution at room temperature. Using a pipette with 10μm of ASW, we transferred a specimen into the same dish and placed it on ice for about 10sec, which prevented damage to the animals. Thus, we embedded a placozoan in the middle of the agar drop, and a small agar block with the animal was placed into a recording chamber filled with ASW or a given extracellular solution. The small sizes of agar block hold the animals, allowed to perform microelectrode recordings and change solutions as needed. Placozoans were monitored under DIC microscopy and were well-maintained for several hours for long-term observations and physiological tests. Statistical analyses of all electrophysiological and pharmacological tests were performed using paired Students’ *t*-tests.

### 2.3. Comparative bioinformatic analyses

We used the data from 18 genomes of basal metazoans and two choanoflagellates (Supplement 1) for the presence of sodium and calcium channels. All accession numbers, gene IDs, and sequences are listed in Supplement 2. The search for possible Na_v_ and Ca_v_ homologs and computational annotation of predicted gene functions was performed using sequence similarity methods(BLAST) algorithm and protein domain detection(Pfam and SMART[43,44]). Human Na_v_ and Ca_v_ protein sequences were used as queries to search target proteomes. All hits with the score at least 100 were put as queries in BLAST search against the SwissProt database to infer their family assignment and completeness.

Protein sequences were aligned in Mafft[45] with default settings. Phylogenetic trees were inferred using the Maximum Likelihood algorithm implemented in IQTREE web server[46]. Tree robustness was tested with 1000 replicates of ultrafast bootstrap.

## 3. Results

### 3.1. Induced action potentials in placozoans

Fig.1 shows one of the tested placozoan species (*Hoilungia*), which are flat animals (20μm in ‘dorso-ventral’ orientation). For electrical recordings, we used extracellular glass microelectrodes with the initial resistance 1-2MΩ. Applying negative pressure inside the pipette on contact with placozoan cells increased the resistance of microelectrodes to 6-18MΩ. In general, no spontaneous action potentials were recorded. However, electrical stimulation (10-150nA pulses, 0.5-2sec) induced a burst of electrical signals (**Fig. 1B,C,H**), and these types of responses were observed in each of the four tested species of Placozoa (**Fig.1S** Supplement). The averaging of 50-1000 electrical responses revealed a class of regenerative electrical signals, which we recognized as action potentials. They were similar to action potentials observed in other basal metazoans and bilaterian animals (including mammals), where recording is performed either from nerves or around neuronal somata[47]. It is also possible to see different shapes of these action potentials (**Figs. 1C,H,2B**), which likely reflect different positions of recording electrodes. Of note, the duration of these action potentials is short (compared to the majority of invertebrates[47]), and the averaging of the signals was ~3ms, which is comparable to some fast-spiking activity (also shorter action potentials, 1-2ms, were recorded).

**Figure 1.**
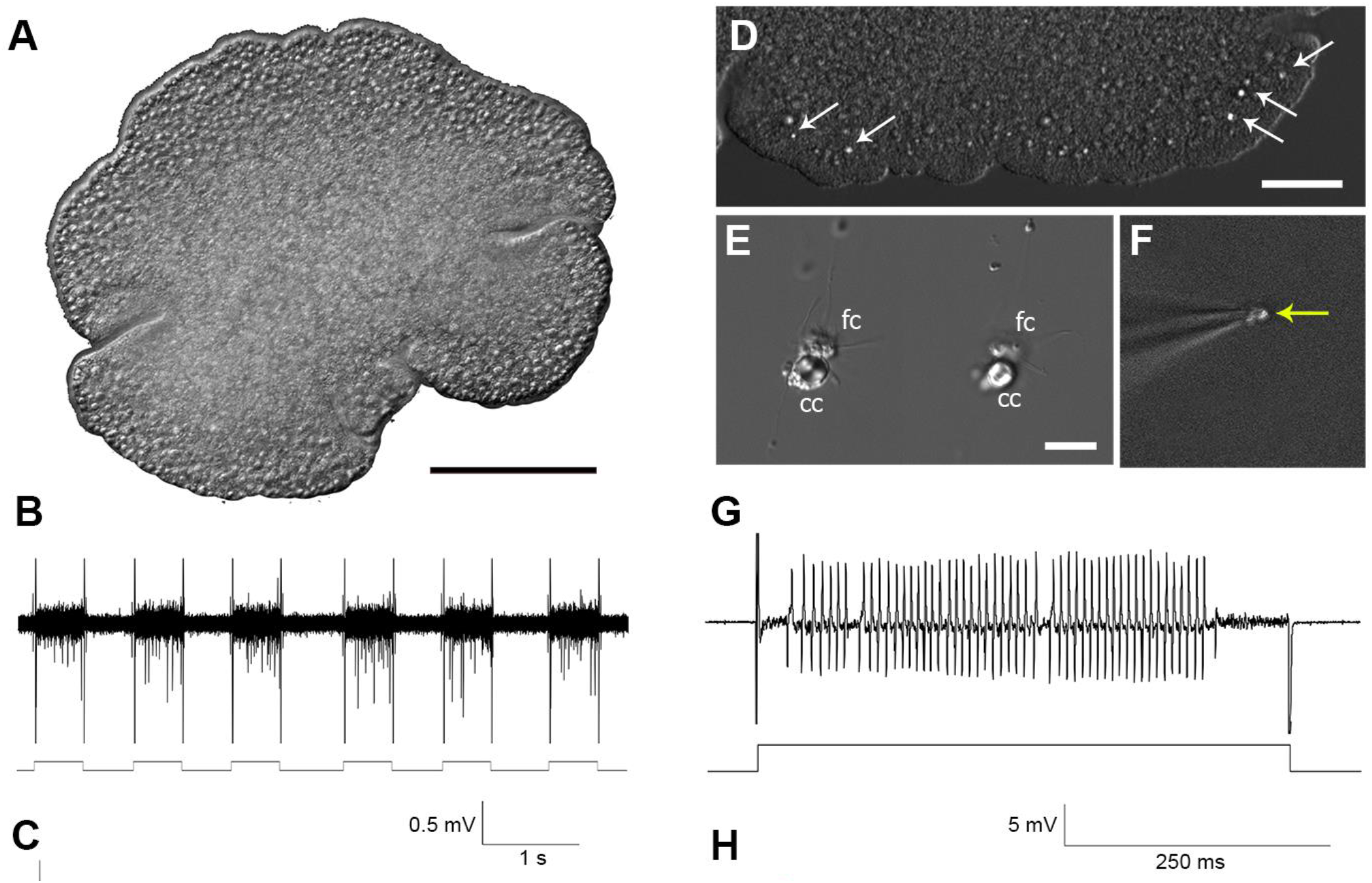
Electrical activity in Placozoa. A) A general view of *Hoilungia hongkongensis* (the focus is on the upper side of this disc-shaped animal). B, C) Extracellular recording from an intact animal. Depolarization by 50nA current repetitively induced a burst of action potentials. C) Illustrative examples of different electrical responses following a single depolarization pulse; an insert shows the shape of a normalized signal, which was obtained by averaging of several dozen action potentials within a given response to a depolarization pulse (lower trace). D-G) Patch recording from a single crystal cell. D) A view of the rim area of *Trichoplax sp*.(H2), arrows show the position of crystal cells (gravity receptors[48]), which can easily be observed under DIC illumination. E) Isolated crystal cells (cc); these cells can be isolated together with fiber cells (fc). F) The image shows the electrode with a single crystal cell (note a visible aragonite crystal - arrow). G) Electrical recording from a single crystal cell (current pulse: 20nA). H) Different shapes of spikes recorded from *Hoilungia*. Scale: A-100μm; D–50μm; E–10 μm.

The specialized crystal cells were recently discovered in both *Trichoplax*[17] and *Hoilungia*[31,41]; they were identified as the gravity sensors[48]. These cells are easily recognized in intact animals under differential contrast illumination due to the presence of a large aragonite crystal in each cell (**Fig. 1D**). These cells are also easy to isolate (**Fig. 1E**), and they are relatively large for single-cell recording (**Fig. 1F**). Fig 1G shows a prominent burst of fast action potentials induced in the crystal cells.

### 3.2. Sodium-dependence of spikes in placozoans

Short durations of the recorded electrical signals in placozoans suggest that these action potentials reflect the activity of sodium-type ion channels. To test this possibility, we replaced the standard artificial seawater with sodium-free solution, were NaCl was substituted by *N*-methyl-d-glucamine (NMDG) to preserve the osmolarity. As in control tests – animals were immobilized in small blocks of 1% agarose, which limited movements of animals but allowed changing of solutions. The induced electrical activity was reversibly eliminated in Na-free NMDG solution, suggesting sodium-dependence of observed action potentials (**Fig.2**). Of note, direct placement of free-moving placozoans in the NMDG solution induced dissociation of animals into cells within 30-60 mins. Thus, as a control test, we placed animals in the 1% agar block made of Na-free solution, where the animals preserved their shape; then, we were able to use an extracellular electrode filled with different solutions to test the presence/absence of Na-dependent action potentials. Applying a small positive pressure during the recording, we carried out microapplication of the solution inside the electrode directly to the recording site. Here, we showed that when the electrode was filled with Na-free solution, no action potentials were induced by current pulses, but the spikes were restored when the electrode filled with NaCl-containing artificial seawater was used (**Fig. 2C,D**;n=14).

**Figure 2.**
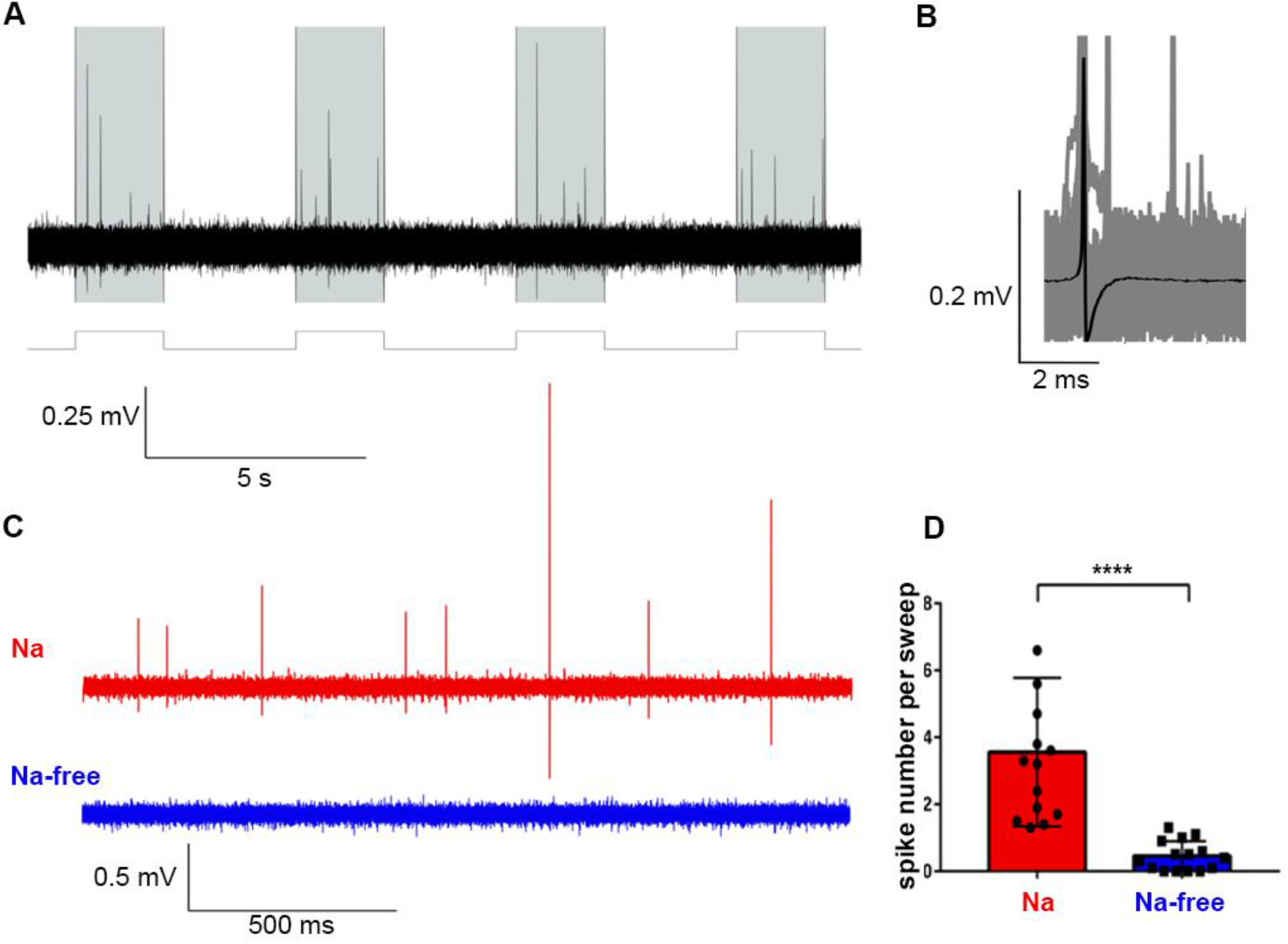
A) Action potentials in response to 150nA current pulses (shaded area, 2s). B) The averaged action potential (AP) from all recorded action potentials (n=911). C) An illustrated example of a typical response to a single current pulse in the artificial seawater [ASW] (with Na^+^, red trace) and in the Na^+^-free solution (NaCl was replaced by N-methyl-D-glucamine (NMDG), blue trace). D) A histogram shows average numbers of APs to single current pulse in the presence of Na^+^ [red,ASW] and in the Na^+^-free solution(blue); Students’ paired *t*-test, ****p<0.0001,n=14).

### 3.3 The Diversity and Structure of sodium channels in Placozoa

Na^+^-dependent action potentials in placozoans could be mediated by two classes of voltage-gated cation channels (Ca_v_ and Na_v_, **Fig.3B**) with a shared topology containing 24 transmembrane (TM) helices, which are four repeats of 6TM domains[36,37,40,49,50] evolved from prokaryotic ancestors[37,51]. One sodium-conducting Ca_v_3 and two Na_v_ channels in *Trichoplax* have been reported[6,36,37,50]. Here, we identified five Na_v_ in both *Trichoplax* species (H1, H2 haplotypes) and seven channels in each species of the genus *Hoilungia* (**Figs. 3A, and 4**). It appears that in the lineage leading to *Hoilungia*, there was one additional duplication event generating novel Na_v_ orthologs (**Fig. 4**). Thus, the discovered diversity of Na_v_-type channels in Placozoa is significantly higher than it was anticipated and observed in other basal metazoans, including ctenophores, sponges, and most of the bilaterians, except vertebrates.

**Figure 3.**
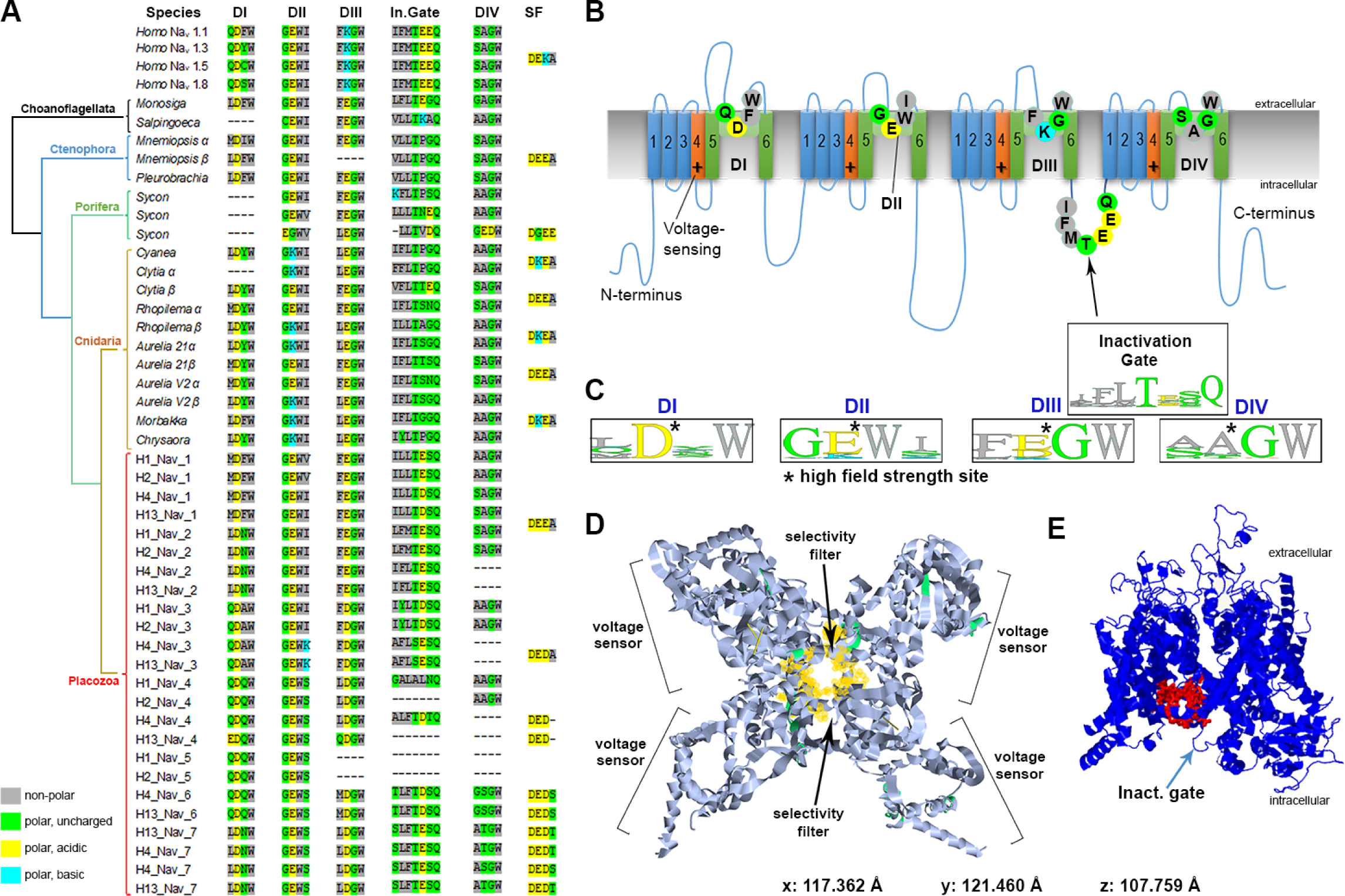
Voltage-gated sodium channels(Na_v_) in Placozoa revealed a remarkable diversity of ion selectivity pore motifs. All metazoan’s Na_v_ have the same 4-domain organization, acquired from ancestral eukaryotes[6,36,38,40,49,55]. Here, choanoflagellates are used as the outgroup (Fig.4). A) The table shows critical amino acids of Na_v_ domains (D1-DIV), which contribute to pore selectivity motifs (marked as SF–<u>S</u>electivity <u>F</u>ilter)[37]) and inactivation gates (In.Gate). Top four entries represent human Na_v_1; these channels contain a critical lysine (K) in the pore region, which is responsible for sodium selectivity[6,36,37]. B) Schematic organization of a generalized sodium channel. Each of the four Na_v_ domain (I-IV) contains six transmembrane loops (1-6), the pore region with four key amino-acids (indicated by circles) responsible for ion selectivity (table A). The voltage sensor is indicated as **+**. A region responsible for inactivation is located between domains DIII and DIV. Selectivity pore motifs are from *Homo* Na*v*1 subtypes[6,36,37]. C) WebLogo representation[56] of four critical amino acids forming ion selectivity filters (SF), as calculated from table A for each of the Na_v_ domains (DI–DIV). D-E) The reconstruction of a hypothetical 3D-structure of the Na_v_-1-like channel from *Trichoplax adhaerens* (TriadITZ_003340). D) Top Na_v_ view is based on pdb ID: 6A90 modeling[57]). E) Side view of the same Na_v_-1-like *Trichoplax* channel generated using Phyre2 modeling[58]. The structural model is close to the human Na_v_1.4 type, where a ‘detection’ pocket is marked by red, and located close to the inactivation gate (arrow).

**Figure 4.**
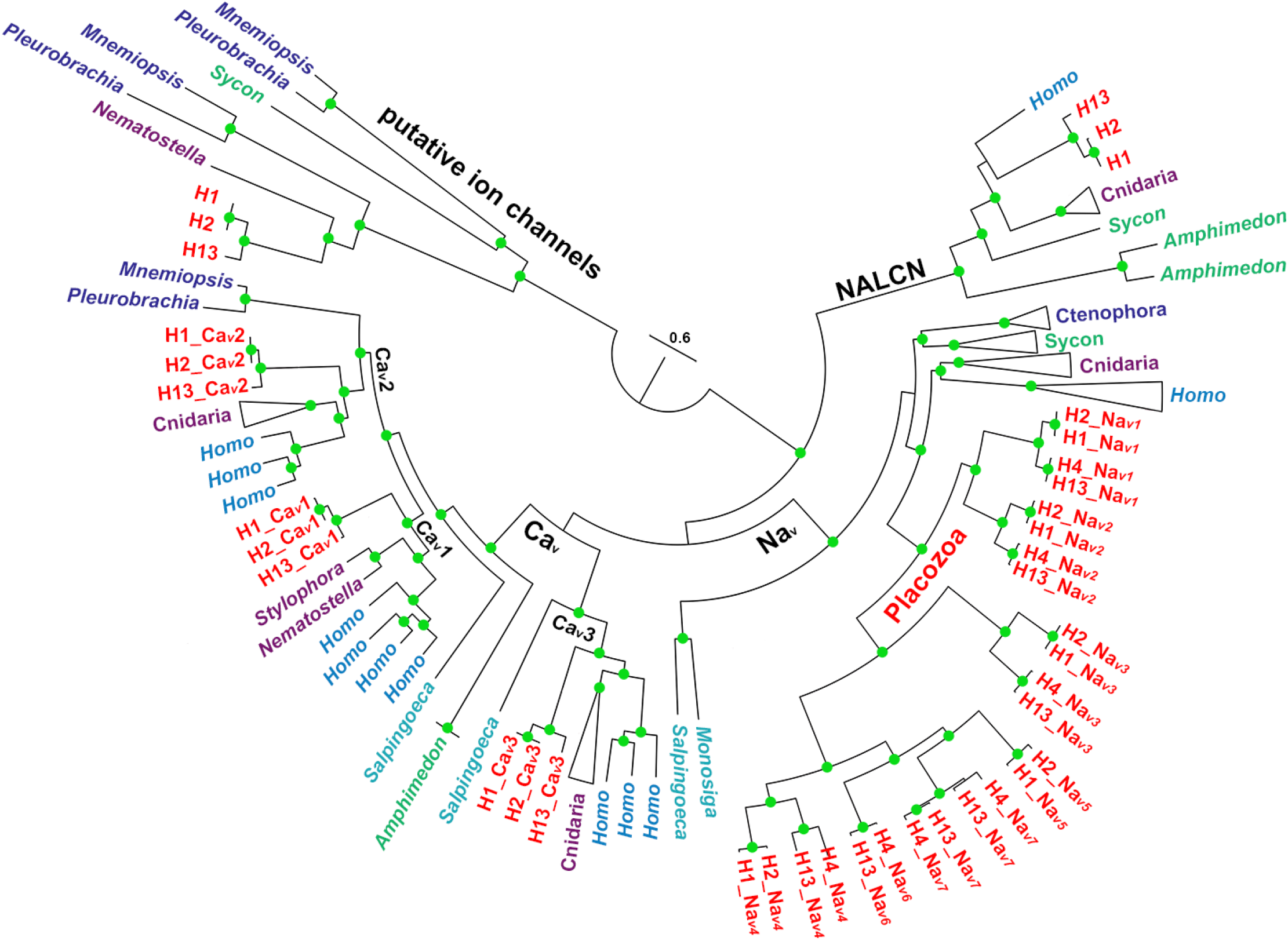
The diversity and evolutionary relationships of voltage-gated sodium and calcium channels in Metazoa and two species of choanoflagellates. (*Monosiga* and *Salpingoeca*) as outgroups. 207 protein sequences were aligned with Mafft[45]. The phylogenetic tree was reconstructed using Iqtree[46] with the VT+G4 evolutionary model. Abbreviations: Na_v_–voltage-gated sodium channels, Ca_v_-voltage-gated calcium channels, NALCN–sodium leak channels. Clade labeled “Putative ion channels” correspond to the group of sequences with no experimentally studied homologs, which mostly present in early-branching metazoan lineages. Placozoan species are denoted by mitochondrial haplotypes: H1–*Trichoplax adhaerens*, H2– *Trichoplax* sp., H13–*Hoilungia hongkongensis*, H4–*Hoilungia* sp. Selective groups of ctenophores, cnidaria, and sponges are collapsed here, but Fig.1S(supplement) shows the expanded version of this tree. The references for each particular species, gene and/or their sequences with relevant GeneBank accession numbers are summarized in the supplementary excel Table S1.

The pore motifs controlling calcium(EEEE/EEDD) or sodium(DEKA/DKEA) ion selectivity is highly conservative over 800 million years of evolution and can be predicted based on the presence of negatively charged amino acids such as aspartate(D) and glutamate(E) and a positively charged lysine(K)[6,36–38,52,53]. **Fig. 3A** illustrates a much greater diversity of the pore/ion selectivity filter motifs discovered in placozoans compared to other studied groups, including cnidarians and many bilaterians (Supplement 3-6).

Of note, some pore motifs contain polar uncharged amino acids - threonine(T) and serine (S), suggesting distinct characteristics of voltage-gated channels in Placozoa. Specifically, in both *Trichoplax* and *Hoilungia*, we identified novel combinations for selective filters responsible for Na^+^ pore permeability (**Fig.3A**): DE<u>D</u>A, DEE<u>T</u>, DE<u>DT</u>, DE<u>DS</u>, and already known from other species DEEA, characteristic for Na_v_2-type of the channels with less specific cation selectivity[37].

## Discussion

The observed diversity of Na_v_ channels and their motifs in Placozoa also implies differences in their functions, cellular and subcellular localization, and pore selectivity – all to be tested experimentally in future studies. The presence of distinct pore motifs suggests that both Na^+^ and Ca^2+^ permeability occurred in Na_v_-like and Ca_v_3-like channels[34,37,50]. The involvement of Ca_v_3-like channels in Na-dependent action potentials is highly likely, but their kinetics can be slower[34].

Some medusozoans (Cnidaria) and bilaterians (including humans) independently developed Na_v_1-type channels with very fast activation kinetics and high selectivity for Na^+^ vs. Ca^2+^[6,36,37,40,49,54]. This metazoan-specific innovation is associated with the presence of a single positively-charged lysine (K) in the selective filter region, leading to DE<u>K</u>A and D<u>K</u>EA motifs for humans and jellyfishes, respectively[36,37,40,49,51]. In two species of *Hoilungia* (H4 and H13 haplotypes), we also identified lysine (K) but in the 4th vs. the 2nd positions (as in jellyfishes) of the second domain (DII, **Fig. 3A**), suggesting faster kinetics for placozoans’ Na_v_.

We detect genes encoding Na_v_-type channels in ctenophores (which is not surprising due to their predatory lifestyle and complex behaviors) and in the sessile calcareous sponge *Sycon*; Fig. 4 shows that additional/putative voltage-gated cationic channels can be discovered in basal metazoans (although we do not exclude the presence of possible pseudogenes). In contrast, the desmosponges likely lost sodium voltage-gated channels.

The systemic functions of ion channels in placozoans are unknown. The detection of the action potential in the crystal cells is an important indication of the role of fast signaling in the control of the geotaxis and spatial orientation. However, systematic analysis of cellular and subcellular expression of Na_v_ as well as pharmacological and imaging studies would be essential to characterize cellular and biophysical bases of rapid (Na-dependent) responses in these animals. The diversity of predicted Na_v_ channels might also reflect the resistance of placozoans against potential toxins.

In summary, the revealed adaptive radiation of Na_v_ in Placozoa and the appearance of additional channel orthologs in *Hoilungia* (compared to *Trichoplax*) emphasize the importance of Placozoa for deciphering the evolution of integrative functions in animals and discoveries of novel properties of ion channels in these cryptic animals.

## Acknowledgments

This work was supported, in part, by the Human Frontiers Science Program (RGP0060/2017) and National Science Foundation (1146575,1557923,1548121 and 1645219) grants to L.L.M.; and Russian Ministry of Science and High Education (2020-1902-01-312) grant to P.M.B.

## Role of authors

All authors had access to the data in the study and took responsibility for the integrity of the data and the accuracy of the data analysis. DYR, IVS and MAN share authorship equally. Research design: LLM and DYR. Molecular data and sequencing analyses: DYR, MAN, ABK, LLM; Phylogeny: MAN. Electrophysiological experiments: IVS and DYR. Analysis and interpretation of data: all authors. Drafting of the article: DYR and LLM.

## References

[1] P.A. Anderson, R.M. Greenberg, Phylogeny of ion channels: clues to structure and function, Comp Biochem Physiol B Biochem Mol Biol 129 (2001) 17–28. 10.1016/s1096-4959(01)00376-1.

[2] X. Cai, Ancient origin of four-domain voltage-gated Na+ channels predates the divergence of animals and fungi, J Membr Biol 245 (2012) 117–123. 10.1007/s00232-012-9415-9.

[3] A.M. Castelfranco, D.K. Hartline, Evolution of rapid nerve conduction, Brain Res 1641 (2016) 11–33. 10.1016/j.brainres.2016.02.015.

[4] L.L. Moroz, K.M. Kocot, M.R. Citarella, S. Dosung, T.P. Norekian, I.S. Povolotskaya, A.P. Grigorenko, C. Dailey, E. Berezikov, K.M. Buckley, A. Ptitsyn, D. Reshetov, K. Mukherjee, T.P. Moroz, Y. Bobkova, F. Yu, V.V. Kapitonov, J. Jurka, Y.V. Bobkov, J.J. Swore, D.O. Girardo, A. Fodor, F. Gusev, R. Sanford, R. Bruders, E. Kittler, C.E. Mills, J.P. Rast, R. Derelle, V.V. Solovyev, F.A. Kondrashov, B.J. Swalla, J.V. Sweedler, E.I. Rogaev, K.M. Halanych, A.B. Kohn, The ctenophore genome and the evolutionary origins of neural systems, Nature 510 (2014) 109–114. 10.1038/nature13400.

[5] L.L. Moroz, A.B. Kohn, Independent origins of neurons and synapses: insights from ctenophores, Philos Trans R Soc Lond B Biol Sci 371 (2016) 20150041. 10.1098/rstb.2015.0041.

[6] B.J. Liebeskind, D.M. Hillis, H.H. Zakon, Evolution of sodium channels predates the origin of nervous systems in animals, Proc Natl Acad Sci U S A 108 (2011) 9154–9159. 10.1073/pnas.1106363108.

[7] M. Srivastava, O. Simakov, J. Chapman, B. Fahey, M.E. Gauthier, T. Mitros, G.S. Richards, C. Conaco, M. Dacre, U. Hellsten, C. Larroux, N.H. Putnam, M. Stanke, M. Adamska, A. Darling, S.M. Degnan, T.H. Oakley, D.C. Plachetzki, Y. Zhai, M. Adamski, A. Calcino, S.F. Cummins, D.M. Goodstein, C. Harris, D.J. Jackson, S.P. Leys, S. Shu, B.J. Woodcroft, M. Vervoort, K.S. Kosik, G. Manning, B.M. Degnan, D.S. Rokhsar, The Amphimedon queenslandica genome and the evolution of animal complexity, Nature 466 (2010) 720–726. 10.1038/nature09201.

[8] S.P. Leys, G.O. Mackie, R.W. Meech, Impulse conduction in a sponge, J Exp Biol 202 (Pt 9) (1999) 1139–1150.

[9] S.P. Leys, G.O. Mackie, Electrical recording from a glass sponge, Nature 387 (1997) 29–30. doi.org/10.1038/387029b0.

[10] T. Monk, M.G. Paulin, Predation and the origin of neurones, Brain Behav Evol 84 (2014) 246–261. 10.1159/000368177.

[11] M.J. Telford, L.L. Moroz, K.M. Halanych, Evolution: A sisterly dispute, Nature 529 (2016) 286–287. 10.1038/529286a.

[12] L.L. Moroz, NeuroSystematics and Periodic System of Neurons: Model vs Reference Species at Single-Cell Resolution, ACS Chem Neurosci 9 (2018) 1884–1903. 10.1021/acschemneuro.8b00100.

[13] F. Varoqueaux, D. Fasshauer, Getting Nervous: An Evolutionary Overhaul for Communication, Annu Rev Genet 51 (2017) 455–476. 10.1146/annurev-genet-120116-024648.

[14] M. Srivastava, E. Begovic, J. Chapman, N.H. Putnam, U. Hellsten, T. Kawashima, A. Kuo, T. Mitros, A. Salamov, M.L. Carpenter, A.Y. Signorovitch, M.A. Moreno, K. Kamm, J. Grimwood, J. Schmutz, H. Shapiro, I.V. Grigoriev, L.W. Buss, B. Schierwater, S.L. Dellaporta, D.S. Rokhsar, The *Trichoplax* genome and the nature of placozoans, Nature 454 (2008) 955–960. 10.1038/nature07191.

[15] B. Schierwater, R. DeSalle. Placozoa, Curr Biol 28 (2018) R97–R98. 10.1016/j.cub.2017.11.042.

[16] K.G. Grell, A. Ruthmann Placozoa, in: F.W. Harrison (Ed.) Microscopic Anatomy of Invertebrates, Wiley-Liss, New York, 1991, pp. 13–27.

[17] C.L. Smith, F. Varoqueaux, M. Kittelmann, R.N. Azzam, B. Cooper, C.A. Winters, M. Eitel, D. Fasshauer, T.S. Reese, Novel.cel. types, n.urosecretor. cells, and body plan of the early-diverging metazoan *Trichoplax adhaerens*, Curr Biol 24 (2014) 1565–1572. 10.1016/j.cub.2014.05.046.

[18] A. Senatore, T.S. Reese, C.L. Smith, Neuropeptidergic integration of behavior in *Trichoplax adhaerens*, an animal without synapses, J Exp Biol 220 (2017) 3381–3390. 10.1242/jeb.162396.

[19] C.L. Smith, N. Pivovarova, T.S. Reese, Coordinated Feeding Behavior in *Trichoplax*, an Animal without Synapses, PLoS One 10 (2015) e0136098. 10.1371/journal.pone.0136098.

[20] T. Ueda, S. Koya, Y.K. Maruyama, Dynamic patterns in the locomotion and feeding behaviors by the placozoan *Trichoplax adhaerence*., Biosystems 54 (1999) 65–70.

[21] C.L. Smith, T.S. Reese, T. Govezensky, R.A. Barrio, Coherent directed movement toward food modeled in *Trichoplax*, a ciliated animal lacking a nervous system, Proc Natl Acad Sci U S A 116 (2019) 8901–8908. 10.1073/pnas.1815655116.

[22] D.Y. Romanova, A. Heyland, D. Sohn, A.B. Kohn, D. Fasshauer, F. Varoqueaux, L.L. Moroz, Glycine as a signaling molecule and chemoattractant in *Trichoplax* (Placozoa): insights into the early evolution of neurotransmitters, Neuroreport 31 (2020) 490–497. 10.1097/WNR.0000000000001436.

[23] A. Heyland, R. Croll, S. Goodall, J. Kranyak, R. Wyeth, *Trichoplax adhaerens*, an enigmatic basal metazoan with potential, Methods Mol Biol 1128 (2014) 45–61. 10.1007/978-1-62703-974-1_4.

[24] A. Fortunato, A. Aktipis, Social feeding behavior of *Trichoplax adhaerens*, Front Ecol Evol 7 (2019). 10.3389/fevo.2019.00019.

[25] S. Armon, M.S. Bull, A. Aranda-Diaz, M. Prakash, Ultrafast epithelial contractions provide insights into contraction speed limits and tissue integrity, Proc Natl Acad Sci U S A 115 (2018) E10333–E10341. 10.1073/pnas.1802934115.

[26] T.D. Mayorova, K. Hammar, C.A. Winters, T.S. Reese, C.L. Smith, The ventral epithelium of *Trichoplax adhaerens* deploys in distinct patterns cells that secrete.digestiv. enzymes, mucus or diverse neuropeptides, Biol Open 8 (2019). 10.1242/bio.045674.

[27] L.L. Moroz, D.Y. Romanova, M.A. Nikitin, D. Sohn, A.B. Kohn, E. Neveu, F. Varoqueaux, D. Fasshauer, The diversification and lineage-specific expansion of nitric oxide signaling in Placozoa: insights in the evolution of gaseous transmission, Sci Rep 10 (2020) 13020. 10.1038/s41598-020-69851-w.

[28] L.L. Moroz, D. Sohn, D.Y. Romanova, A.B. Kohn, Microchemical identification of enantiomers in early-branching animals: Lineage-specific diversification in the usage of D-glutamate and D-aspartate, Biochem Biophys Res Commun 527 (2020) 947–952. 10.1016/j.bbrc.2020.04.135.

[29] M. Nikitin, Bioinformatic prediction of *Trichoplax adhaerens* regulatory peptides, Gen Comp Endocrinol 212 (2015) 145–155. 10.1016/j.ygcen.2014.03.049.

[30] F. Varoqueaux, E.A. Williams, S. Grandemange, L. Truscello, K. Kamm, B. Schierwater, G. Jekely, D. Fasshauer, High Cell Diversity and Complex Peptidergic Signaling Underlie Placozoan Behavior, Curr Biol 28 (2018) 3495–3501 e3492. 10.1016/j.cub.2018.08.067.

[31] M. Eitel, W.R. Francis, F. Varoqueaux, J. Daraspe, H.J. Osigus, S. Krebs, S. Vargas, H. Blum, G.A. Williams, B. Schierwater, G. Worheide, Comparative genomics and the nature of placozoan species, PLoS Biol 16 (2018) e2005359. 10.1371/journal.pbio.2005359.

[32] H.J. Osigus, S. Rolfes, R. Herzog, K. Kamm, B. Schierwater, *Polyplacotoma mediterranea* is a new ramified placozoan species, Curr Biol 29 (2019) R148–R149. 10.1016/j.cub.2019.01.068.

[33] C.L. Smith, T.S. Reese, Adherens Junctions Modulate Diffusion between Epithelial Cells in *Trichoplax adhaerens*, Biol Bull 231 (2016) 216–224. 10.1086/691069.

[34] C.L. Smith, S. Abdallah, Y.Y. Wong, P. Le, A.N. Harracksingh, L. Artinian, A.N. Tamvacakis, V. Rehder, T.S. Reese, A. Senatore, Evolutionary insights into T-type Ca(2+) channel.structure. function, and ion selectivity from the *Trichoplax adhaerens* homologue, J Gen Physiol 149 (2017) 483–510. 10.1085/jgp.201611683.

[35] W. Elkhatib, C.L. Smith, A. Senatore, A Na(+) leak channel cloned from *Trichoplax adhaerens* extends extracellular pH and Ca(2+) sensing for the DEG/ENaC family close to the base of Metazoa, J Biol Chem 294 (2019) 16320–16336. 10.1074/jbc.RA119.010542.

[36] H.H. Zakon, Adaptive evolution of voltage-gated sodium channels: the first 800 million years, Proc Natl Acad Sci U S A 109 Suppl 1 (2012) 10619–10625. 10.1073/pnas.1201884109.

[37] J.E. Fux, A. Mehta, J. Moffat, J.D. Spafford, Eukaryotic Voltage-Gated Sodium Channels: On.Thei. Origins, A.ymmetries. Losses, Diversification and Adaptations, Front Physiol 9 (2018) 1406. 10.3389/fphys.2018.01406.

[38] Y. Moran, M.G. Barzilai, B.J. Liebeskind, H.H. Zakon, Evolution of voltage-gated ion channels at the emergence of Metazoa, J Exp Biol 218 (2015) 515–525. 10.1242/jeb.110270.

[39] A. Ghezzi, B.J. Liebeskind, A. Thompson, N.S. Atkinson, H.H. Zakon, Ancient association between cation leak channels and Mid1 proteins is conserved in fungi and animals, Front Mol Neurosci 7 (2014) 15. 10.3389/fnmol.2014.00015.

[40] B.J. Liebeskind, D.M. Hillis, H.H. Zakon, Independent acquisition of sodium selectivity in bacterial and animal sodium channels, Curr Biol 23 (2013) R948–949. 10.1016/j.cub.2013.09.025.

[41] D.Y. Romanova, Cell types diversity of H4 haplotype Placozoa sp., Marine Biological Journal 4 (2019) 81–90. 10.21072/mbj.2019.04.1.07.

[42] J.B. Thuma, S.L. Hooper, Choline and NMDG directly reduce outward currents: reduced outward current when these substances replace Na(+) is alone not evidence of Na(+)-activated K(+) currents, J Neurophysiol 120 (2018) 3217–3233. 10.1152/jn.00871.2017.

[43] T.C. Giles, R.D. Emes, Inferring Function from Homology, Methods Mol Biol 1526 (2017) 23–40. 10.1007/978-1-4939-6613-4_2.

[44] I. Letunic, P. Bork, 20 years of the SMART protein domain annotation resource, Nucleic Acids Res 46 (2018) D493–D496. 10.1093/nar/gkx922.

[45] K. Katoh, D.M. Standley, MAFFT multiple sequence alignment software version 7: improvements in performance and usability, Mol Biol Evol 30 (2013) 772–780. 10.1093/molbev/mst010.

[46] J. Trifinopoulos, L.T. Nguyen, A. von Haeseler, B.Q. Minh, W-IQ-TREE: a fast online phylogenetic tool for maximum likelihood analysis, Nucleic Acids Res 44 (2016) W232–235. 10.1093/nar/gkw256.

[47] T.H. Bullock, G.A. Horridge, Structure and Function in the Nervous Systems of Invertebrates., Freeman, San Francisco, 1965.

[48] T.D. Mayorova, C.L. Smith, K. Hammar, C.A. Winters, N.B. Pivovarova, M.A. Aronova, R.D. Leapman, T.S. Reese, Cells containing aragonite crystals mediate responses to gravity in *Trichoplax adhaerens* (Placozoa), an animal lacking neurons and synapses, PLoS One 13 (2018) e0190905. 10.1371/journal.pone.0190905.

[49] B.J. Liebeskind, D.M. Hillis, H.H. Zakon, Convergence of ion channel genome content in early animal evolution, Proc Natl Acad Sci U S A 112 (2015) E846–851. 10.1073/pnas.1501195112.

[50] A. Senatore, H. Raiss, P. Le, Physiology and Evolution of Voltage-Gated Calcium Channels in Early Diverging Animal.Phyla.Cnidaria, Placozoa, Porifera and Ctenophora, Front Physiol 7 (2016) 481. 10.3389/fphys.2016.00481.

[51] A. Nishino, Y. Okamura, Evolutionary History of Voltage-Gated Sodium Channels, Handb Exp Pharmacol 246 (2018) 3–32. 10.1007/164_2017_70.

[52] H.H. Zakon, M.C. Jost, Y. Lu, Expansion of voltage-dependent Na+ channel gene family in early tetrapods coincided with the emergence of terrestriality and increased brain complexity, Mol Biol Evol 28 (2011) 1415–1424. 10.1093/molbev/msq325.

[53] I. Pozdnyakov, O. Matantseva, S. Skarlato, Diversity and evolution of four-domain voltage-gated cation channels of eukaryotes and their ancestral functional determinants, Sci Rep 8 (2018) 3539. 10.1038/s41598-018-21897-7.

[54] F.H. Yu, W.A. Catterall, Overview of the voltage-gated sodium channel family, Genome Biol 4 (2003) 207. 10.1186/gb-2003-4-3-207.

[55] B.J. Liebeskind, D.M. Hillis, H.H. Zakon, Phylogeny unites animal sodium leak channels with fungal calcium channels in.a. ancient, voltage-insensitive clade, Mol Biol Evol 29 (2012) 3613–3616 10.1093/molbev/mss182.

[56] G.E. Crooks, G. Hon, J.M. Chandonia, S.E. Brenner, WebLogo: a sequence logo generator, Genome Res 14 (2004) 1188–1190. 10.1101/gr.849004.

[57] H. Berman, K. Henrick, H. Nakamura, Announcing the worldwide Protein Data Bank, Nat Struct Biol 10 (2003) 980. 10.1038/nsb1203-980.

[58] L.A. Kelley, S. Mezulis, C.M. Yates, M.N. Wass, M.J. Sternberg, The Phyre2 web portal for.protei. modeling, prediction and analysis, Nat Protoc 10 (2015) 845–858. 10.1038/nprot.2015.053.

